# Exploring the functional, protective, and transcriptomic effects of GIP on cytokine-exposed human pancreatic islets and EndoC-βH5 cells

**DOI:** 10.1101/2024.06.17.599254

**Authors:** Kristine Henriksen, Anne Jørgensen, Simranjeet Kaur, Rebekka Gerwig, Cecilie Amalie Brøgger Svane, Filip K. Knop, Joachim Størling

**Affiliations:** Translational Type 1 Diabetes Research, Department of Clinical and Translational Research, Steno Diabetes Center Copenhagen,, Denmark; Department of Biomedical Sciences, Faculty of Health and Medical Sciences, University of Copenhagen, Copenhagen, Denmark; Center for Clinical Metabolic Research, Gentofte Hospital, University of Copenhagen, Hellerup, Denmark; Department of Clinical Medicine, Faculty of Health and Medical Sciences, University of Copenhagen, Copenhagen, Denmark; Translational Type 2 Diabetes Research, Department of Clinical and Translational Research, Steno Diabetes Center Copenhagen, Herlev, Denmark

**Author notes:** **Correspondence:** Joachim Størling, MSc Ph.D, Translational Omics and Islet Biology, Translational Type 1 Diabetes Research, Clinical and Translational Research, Steno Diabetes Center Copenhagen, Borgmester Ib Juuls Vej 83, 2730 Herlev, Denmark. **Currently employed at:** Novo Nordisk A/S, Søborg, Denmark (Filip K. Knop).

**Keywords:** GIP, glucose-dependent insulinotropic polypeptide, type 1 diabetes, human islets, EndoC-βH5, proinflammatory cytokines, IL-1β, IFN-y, apoptosis

## Abstract

Immune-mediated beta-cell destruction and lack of alpha-cell responsiveness to hypoglycaemia are hallmarks of type 1 diabetes pathology. The incretin hormone glucose-dependent insulinotropic polypeptide (GIP) may hold therapeutic potential for type 1 diabetes due to its insulinotropic and glucagonotropic effects, as well as its beta-cell protective effects shown in rodent islets. Here, we examined the functional, protective, and transcriptomic effects of GIP treatment upon diabetogenic cytokine exposure to interleukin (IL)-1β ± interferon (IFN)-γ in human EndoC-βH5 beta cells and isolated human islets, respectively.

GIP dose-dependently augmented glucose-stimulated insulin secretion from EndoC-βH5 cells and increased insulin and glucagon secretion from human islets during high and low glucose concentrations, respectively. The insulinotropic effect of GIP in EndoC-βH5 cells was abrogated by KN-93, an inhibitor of calcium/calmodulin-dependent protein kinase 2 (CaMK2). GIP did not prevent cytokine-induced apoptosis or cytokine-induced functional impairment of human EndoC-βH5 cells. GIP also did not prevent cytokine-induced apoptosis in human islets. GIP treatment of human islets with or without cytokines for 24 hours did not significantly impact the transcriptome. GIP potentiated cytokine-induced secretion of IL-10 and c-c motif chemokine ligand (CCL)-2 from human islets while decreasing the secretion of c-x-c motif chemokine ligand (CXCL)-8. In EndoC-βH5 cells, GIP reduced IFN-γ-induced secretion of tumor necrosis factor (TNF)-α, IL-2, IL-6, and IL-10 but increased the secretion of CXCL8, CCL2, CCL4, and CCL11.

In conclusion, our results suggest that the insulinotropic effect of GIP is CaMK2-dependent. Furthermore, our results indicate that GIP does not provide substantial cytoprotective effects against diabetogenic cytokine challenge or significantly modulate the transcriptome of human islets when applied at a supraphysiological level. GIP may, however, still exert selective inflammation-modulatory effects upon diabetogenic cytokine exposure.

## 1. Introduction

Type 1 diabetes is characterised by immune-mediated and inflammatory destruction of insulin-producing beta cells, leading to hyperglycaemia and the need for life-long insulin replacement therapy. In addition, type 1 diabetes individuals are at a heightened risk of iatrogenic hypoglycaemia due to an impaired counterregulatory glucagon response [1–4]. Despite improvements in insulin replacement therapy, there is a strong demand for adjunctive therapies that mitigate both hyperglycaemia and hypoglycaemia in type 1 diabetes.

In type 1 diabetes, the local inflammatory milieu in the islets of Langerhans is in part driven by the production of proinflammatory cytokines, such as interleukin-1 beta (IL-1β) and interferon-gamma (IFN-γ) [5]. Interestingly, cytokines stimulate both alpha and beta cells to produce and secrete cytokines and chemokines, exacerbating islet inflammation and immune cell infiltration [6–8].

The incretin hormone glucose-dependent insulinotropic polypeptide (GIP) plays a key role in glucose homeostasis with bi-directional effects that contribute to the maintenance of glycaemic control [9, 10]. Upon nutrient intake, GIP is secreted by enteroendocrine K cells and acts on pancreatic beta cells to potentiate glucose-stimulated insulin secretion, promoting glucose deposition to lower and normalise blood glucose [10, 11]. At low plasma glucose concentrations, GIP can stimulate glucagon secretion from pancreatic alpha cells, promoting glucose mobilisation to raise and normalise blood glucose [9, 12–14]. Interestingly, studies in cell lines and rodent models have suggested pro-survival and anti-apoptotic properties of GIP in beta cells [15–18]. Additionally, GIP exhibits complex inflammation-regulatory effects, with results varying based on the model, tissue, and cell type studied yet consistently suggesting an immunomodulatory role [19–22]. Jointly, these putative effects could indicate that GIP holds a therapeutic potential to improve glycaemic control in type 1 diabetes by 1) potentiating insulin secretion from residual beta cells, 2) restoring the counterregulatory glucagon response to hypoglycaemia, and 3) preserving residual beta-cell function and mass via both direct anti-apoptotic effects and via immunomodulatory effects. Nevertheless, most studies investigating the effects of GIP on islet function and survival have been conducted in cell lines and rodent islets. Important species differences in the GIP system limit the translational value of such studies [23–25], necessitating studies in human models to accurately elucidate the therapeutic potential of GIP.

Here, we have investigated the functional, anti-apoptotic, transcriptomic, and immunomodulatory impact of exogenous GIP on human islet cells using two model systems: the non-proliferative human beta-cell line EndoC-βH5 cells [26, 27] and isolated human islets.

## 2. Materials and methods

### 2.1 EndoC-βH5 and human pancreatic islet culture

Frozen vials of ready-to-use EndoC-βH5 cells, a non-proliferative human beta-cell line [26], were seeded in βCoat-coated plates and maintained in Ulti-β1 medium (all from Human Cell Design, France). Isolated human pancreatic islets from non-diabetic and anonymised deceased donors (n = 16, sex (M/F) 9/7, age 42.5 ± 13.9 years, BMI 26.7 ± 3.9 kg/m^2^, HbA1c 5.4 ± 0.3%) were obtained from Prodo Laboratories Inc. (USA) via Tebu-Bio (France). Donor details are summarised in Table 1. Human islets were maintained in F-10 Nutrient Mix + GlutaMAX medium, supplemented with 10% heat-inactivated fetal bovine serum (FBS), 100 U/mL penicillin, and 100 μg/mL streptomycin (all from Gibco, USA). All cells and islets were kept in a humidified incubator at 37°C with 5% CO_2_, and the media was renewed every 2-3 days.

**Table 1.** Islet donor details. Based on the ‘checklist for reporting human islet preparations used in research’ adapted from [74]

| Islet preparation | 1 | 2 | 3 | 4 | 5 | 6 | 7 | 8 |
| --- | --- | --- | --- | --- | --- | --- | --- | --- |
| Donor islets used for | Glucose-dependent secretion assays |  |  |  |  |  |  |  |
| Unique identifier | HP-19136-01 | HP-19150-01 | HP-19163-01 | HP-19170-01 | HP-20296-01 | HP-20304-01 | HP-20317-01 | HP-21013-01 |
| Donor age (years) | 34 | 38 | 24 | 47 | 47 | 45 | 58 | 20 |
| Donor sex (M/F) | M | F | M | M | M | F | M | M |
| Donor BMI (kg/m<sup>2</sup>) | 28.6 | 24.2 | 24.3 | 28.5 | 35.3 | 23.7 | 26.8 | 19.4 |
| Donor HbA<sub>1c</sub> (%) | 5.2 | 5.8 | 5.6 | 5.8 | 5.5 | 4.9 | 4.9 | 5.4 |
| Source of islets | Prodo Laboratories |  |  |  |  |  |  |  |
| Islet isolation centre |  |  |  |  |  |  |  |  |
| History of diabetes? | No | No | No | No | No | No | No | No |

| Islet preparation | 9 | 10 | 11 | 12 | 13 | 14 | 15 | 16 |
| --- | --- | --- | --- | --- | --- | --- | --- | --- |
| Donor islets used for | RNA-sequencing, qPCR and cytokine/chemokine measurements |  |  |  | Cell death assays |  |  |  |
| Unique identifier | HP-19256-01 | HP-19289-01 | HP-19305-01 | HP-19311-01 | HP-21035-01 | HP-21055-01 | HP-21197-01 | HP-21203-01 |
| Donor age (years) | 64 | 51 | 44 | 29 | 65 | 36 | 53 | 25 |
| Donor sex (M/F) | M | F | F | F | F | F | M | M |
| Donor BMI (kg/m<sup>2</sup>) | 25.1 | 24.9 | 24.3 | 26.1 | 25.1 | 31.6 | 32.3 | 26.8 |
| Donor HbA<sub>1c</sub> (%) | 5.4 | 5.2 | 5.8 | 5.1 | 5.2 | 5.6 | 5.5 | 5.8 |
| Source of islets | Prodo Laboratories |  |  |  |  |  |  |  |
| Islet isolation centre |  |  |  |  |  |  |  |  |
| History of diabetes? | No | No | No | No | No | No | No | No |

Abbreviations: F, female; M, Male; BMI, Body mass index; HbA<sub>1c</sub>, hemoglobin A1c;

### 2.2 EndoC-βH5 glucose-simulated insulin secretion

Glucose-stimulated insulin secretion (GSIS) was used to evaluate beta-cell function of EndoC-βH5 cells. Twenty-four hours before the start of the assay, the culture medium was replaced with Ulti-ST starvation medium (Human Cell Design, France). Briefly, cells were washed twice and equilibrated in βKrebs buffer (+ 0.1% fatty acid-free bovine serum albumin) supplemented with 2 mM glucose for 1 hour, followed by incubation for 40 min in βKrebs buffer with either 2 or 20 mM glucose. Collected supernatant samples were analysed for secreted insulin using Human Insulin ELISA (Mercodia, Sweden) according to the manufacturer’s instructions, and luminescence was recorded on the Infinite M200 PRO microplate reader (Tecan, Switzerland). Several experimental setups were used in this study: First, the impact of GIP (0.025-1000 nM, Tocris Bioscience, UK) was evaluated when administered acutely during the GSIS assay. The acute effect of GIP on GSIS was also assessed in the presence of a selection of intracellular signalling inhibitors; H89 (a protein kinase A (PKA) inhibitor, Tocris Bioscience, UK), Bisindolylmaleimide I (a protein kinase C (PKC) inhibitor, Sigma-Aldrich, USA), Nimodipine (an L-type calcium channel (LTCC) inhibitor, Sigma-Aldrich, USA), KN-93 (a calcium/calmodulin-dependent protein kinase II (CaMK2) inhibitor, Sigma-Aldrich, USA), Diazoxide (an inhibitor of K_ATP_-channel closure and membrane depolarisation, Sigma-Aldrich, USA) or vehicle control (equivalent dilution of 0.1% DMSO). The inhibitor doses (each at 10 μM) were based on literature and were present during the equilibration step of GSIS and the 40-min incubation with GIP. Lastly, the impact of GIP on GSIS was evaluated when administered during the 48-hour exposure to IFN-γ (1000 U/mL, Peprotech, USA) or after 48-hour IFN-γ exposure.

### 2.3 Assessment of apoptosis and cytotoxicity in EndoC-βH5 cells

Cell death in EndoC-βH5 cells was measured after 96-hour exposure to IFN-γ ± GIP (0.25-1000 nM) via luminescence-based assays and recorded on the Infinite M200 PRO microplate reader (Tecan, Switzerland). The culture medium was replaced with fresh IFN-γ/GIP after 48 hours. Apoptosis was determined as Caspase 3/7 activity and cell death as dead-cell protease activity using the Caspase-Glo 3/7 Assay or CytoTox-Glo Assay, respectively. Both assays were done according to the manufacturer’s instructions (Promega, USA).

### 2.4 Human islet glucose-dependent hormone secretion

Isolated human islets were handpicked under a stereomicroscope for static glucose-dependent insulin and glucagon secretion assays. Briefly, duplicate groups of 20-25 islets were washed twice in Krebs Ringer HEPES Buffer (KRHB; 115 mM NaCl, 4.7 mM KCl, 2.6 mM CaCl_2_, 1.2 mM KH_2_PO_4_, 1.2 mM MgSO_4_·7H_2_O, 5 mM NaHCO_3_, 20 mM HEPES, 2 mg/mL bovine serum albumin) supplemented with 5.5 mM glucose prior to sequential 30 min incubations of KRBH containing low (2 mM) and high (20 mM) glucose with or without 2.5 nM GIP (Tocris Bioscience, UK, except for donor 1-4 where GIP was from Caslo, Denmark). Secreted insulin and glucagon were determined using Human Insulin ELISA and Human Glucagon ELISA kits (Mercodia, Sweden) and normalised to double-stranded DNA (dsDNA) content to account for differences in islet mass (IEQ) using the QuantiFluor dsDNA system (Promega, USA) as per manufacturing instructions.

### 2.5 Human islet apoptosis assessment

Apoptosis in human islets was determined using Cell Death Detection ELISA (Roche, Switzerland) according to the manufacturer’s instructions. In brief, islets were cultured ±GIP (2.5 nM) and ±IL-1β (R&D systems, USA) and IFN-γ (50 U/mL and 1000 U/mL, respectively) for 48 hours before sample collection. Apoptosis was then measured as the presence of cytoplasmic histone-associated DNA fragments in the cytosol by a sandwich-based ELISA with anti–histone–biotin and anti-DNA-peroxidase antibodies on a streptavidin-coated plate. Absorbance was recorded after the addition of peroxidase substrate ABTS.

### 2.6 Human islet RNA sequencing and data handling

Islets from a total of 4 donors (table 1) were cultured in the presence or absence of IL-1β+IFN-γ ± GIP (2.5 nM, Caslo, Denmark) for 24 hours prior to total RNA extraction using Direct-zol™ RNA Miniprep kit (Zymo _Research_, USA). 100-150 islets in technical duplicates were used per condition. Extracted RNA samples all had RIN values of ≥ 8. A non-stranded, polyA-selected (mRNA) library was prepared and sequenced on the DNBSEQ platform (both by BGI-Copenhagen, Denmark), producing 100-bp-long paired-end reads with an average of 46 million reads per sample. The raw sequencing data were processed as follows; base quality control was performed using FastQC [28]. Pre-processing and QC included the removal of low-quality reads and adapter sequences using SOAPnuke software [29]. Reads were aligned using TopHat (v2.1.1) [30] to the GRCh38 (Ensembl version 107) reference genome. Mapped reads were quantified to gene annotation using HTseq-count (v0.9.1) [31]. Differential gene expression (DGE) analyses (pairwise comparisons between two conditions) were carried out with the DESeq2 package [32] in R. Genes with low read count values (<10 reads and expressed in less than 4 samples) were filtered and removed prior to DGE analysis. Results were visualised with the DESeq2, ggplot2, and pheatmap packages [32–34]. Pathway analyses (Gene Ontology (GO) term enrichment) of up-and downregulated genes were performed using ShinyGO v0.80 [35] with all genes considered in the DGE as background.

### 2.7 RT-qPCR

For real-time quantitative polymerase chain reactions (RT-qPCRs), cDNA was synthesised using the iScript cDNA Synthesis Kit (Bio-Rad, USA), followed by RT-qPCRs using TaqMan Assays and TaqMan Gene Expression Master Mix (both Applied Biosystems, USA) on the CFX384 C1000 Thermal cycler (Bio-Rad, USA). Relative gene expression (fold change) levels were determined with the 2^-ΔΔCt^ method normalising to the housekeeping gene *HRPT1* [36]. Each sample was tested in technical duplicate, and no template controls (NTCs) were included to ensure specificity of TaqMan Assays; B-cell lymphoma 2 (*BCL2*), BCL2 like 1 (*BCL2L1* or *BCL-XL*), BCL2 like 11 (*BCL2L11* or *BIM/BIM-EL*), BCL2 associated X (*BAX*), DNA damage-inducible transcript 3 (*DDIT3* or *CHOP*), Chemokine (C-X-C motif) ligand 10 (*CXCL10*), C-C Motif Chemokine Ligand 5 (*CCL5*), and nitric oxide synthase 2 (*NOS2* or *iNOS*).

### 2.8 Chemokine and cytokine measurements

Accumulated secretion of inflammatory factors (tumour necrosis factor (TNF)-α, interleukin (IL) - 2, -4, -6, -10, -13, Chemokine (C-X-C motif) ligand (CXCL) -8, -10, and C-C Motif Chemokine Ligand (CCL) -2, -3, -4, -11, -13, -17, -22, -26) were measured in the medium from RNA-sequencing experiments of human islets and in stimulation medium from EndoC-βH5 GSIS experiments (both 24h) with the V-PLEX Proinflammatory Panel 1 Human Kit and the V-PLEX Chemokine Panel 1 Human Kit on the MESO Quickplex SQ 120 instrument (all from Meso Scale Diagnostics, USA) according to the manufacturer’s instructions. Sample values below the limit of detection (LOD) were imputed using the LOD/√2 method [37]. For the human islet measurements, secretion was normalised to a factor based on the average of all secreted inflammatory factors for each condition in each donor.

### 2.9 Statistical analyses

Statistical analyses of data (except the RNA-seq and GO term enrichment) were performed using GraphPad Prism (v10, USA) and presented as mean ± standard deviation (SD). Comparisons between two sample groups were analysed with paired Student’s *t-*tests, whereas comparisons between more than two sample groups were analysed by paired one-way or two-way ANOVA with Dunnett’s or Šídák’s multiple comparison tests, as indicated in the figure legends. Statistical significance was defined and reported as a *p*-value < 0.05 (\**p*-value < 0.05, \*\**p*-value < 0.01, \*\*\**p*-value < 0.001), and results were non-significant (ns) unless stated.

## 3. Results

### 3.1 GIP augments glucose-stimulated insulin secretion dose-dependently in EndoC-βH5 cells

To confirm an insulinotropic effect of GIP in the newly established EndoC-βH5 human beta-cell model, we first performed GSIS experiments over a range of supraphysiological GIP doses at low (2 mM) and high (20 mM) glucose (Figure 1a). GIP significantly and dose-dependently increased insulin secretion at high glucose, yielding 244%, 360%, and 446% increases with 0.25, 2.5 and 25 nM GIP, respectively (all *p<*0.001). Raising the GIP dose to 1000 nM did not further increase the incretin effect (398%). GIP also significantly increased insulin secretion at low glucose in a dose-dependent manner, although only maximally at a level corresponding to the baseline level of high glucose alone (*p*=0.008 for 0.25 nM GIP and *p*<0.001 for 2.5-100 nM GIP, Figure 1a).

**Figure 1.**
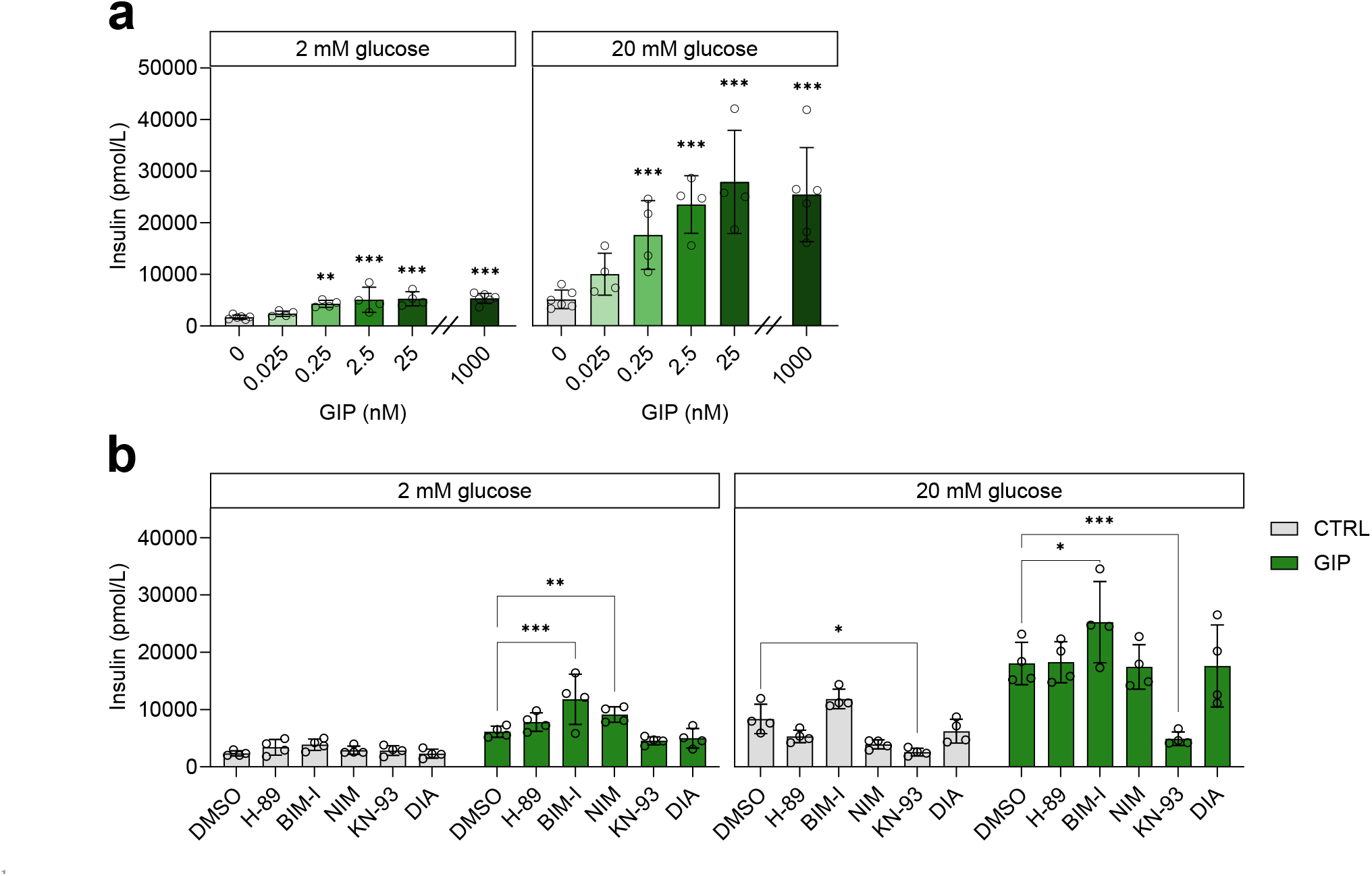
GIP robustly increases glucose-dependent insulin secretion dose-dependently in human EndoC-βH5 cells, which is abrogated by KN-93. (a) Secreted insulin during low glucose (2 mM, left) and during high glucose (20 mM, right) with increasing concentrations (0.025-1000 nM) of GIP. Angled lines on the x-axis denote the transition from x10 increments in GIP dose to 1000 nM GIP. Data are represented as mean ± SD of 4-6 independent experiments. Asterisks (*) indicate statistical significance using mixed-effect analysis/model with Dunnett’s multiple comparisons test each GIP dose compared to 0 nM at 2 or 20 mM glucose, respectively, **p < 0.01, ***p < 0.001. Secreted insulin during low glucose (2 mM, left) and during high glucose (20 mM, right) upon treatment (10 μM each) with H-89 (PKAi), bisindolylmaleimide I (PKCi), nimodipine (LTCCi), KN-93 (CaMK2i), diazoxide (DIA) or vehicle control (0.1% DMSO) without (grey bars) or with GIP (2.5 nM, green bars). Data are represented as mean ± SD of 4 independent experiments. Asterisks (*) indicate statistical significance using paired RM two-way ANOVA with Dunnett’s multiple comparisons test comparing inhibitors to CTRL or GIP vehicle control (DMSO) at 2 or 20 mM glucose, respectively, \**p* <0.05, \*\**p* < 0.01, \*\*\**p* <0.001.

### 3.2 KN-93 abrogates GIP-stimulated insulin secretion in EndoC-βH5 cells

Next, we tested the ability of various small molecule inhibitors of cAMP and calcium signalling to inhibit GIP-induced insulin secretion. We used H-89 (an inhibitor of protein kinase A, PKA), bisindolylmaleimide I (BIM-I; an inhibitor of protein kinase C, PKC), nimodipine (NIM, a blocker of L-type calcium channels, LTCC), KN-93 (an inhibitor of CaMK2), and diazoxide (DIA; an inhibitor of K_ATP_-channel closure). None of the inhibitors altered baseline insulin secretion at low glucose (Figure 1b). However, when co-administered with GIP, BIM-I and NIM increased insulin secretion at low glucose (93%, *p*<0.001 and 49%, *p*=0.009, respectively). At high glucose (Figure 1b), only KN-93 significantly decreased insulin secretion (69%, *p*=0.05). GIP potentiation of insulin secretion at high glucose was unaltered by H-89, NIM and DIA but significantly increased (40%, *p*=0.01) by BIM-I. Remarkably, KN-93 also completely abrogated the insulinotropic effect of GIP at high glucose (*p*=0.007). These results suggest that CaMK2 mediates both glucose-stimulated insulin secretion and the incretin effect of GIP in EndoC-βH5 cells.

### 3.3 GIP does not prevent IFN-γ-induced cell death or functional impairment of EndoC-βH5 cells

We next examined whether GIP could protect against proinflammatory cytokine-induced apoptosis and cytotoxicity. As we have previously established that IFN-γ is the main detrimental diabetogenic cytokine in EndoC-βH5 cells [27], cells were cultured with or without GIP in the presence or absence of IFN-γ for 96 hours (Figure 2). GIP treatment alone neither altered baseline apoptosis evaluated by caspase 3/7 activity (Figure 2a-b) nor baseline cytotoxicity (Figure 2c-d) at any GIP dose tested. IFN-γ exposure induced a 2.14-fold increase in caspase 3/7 activity (*p*=0.03) and a 2.74-fold increase in cytotoxicity (*p*=0.008). GIP treatment alongside IFN-y exposure did not affect IFN-γ-induced apoptosis or cytotoxicity. Since impaired insulin secretion precedes IFN-γ-induced cell death [27], we cultured EndoC-βH5 cells with or without 2.5 nM GIP in the presence or absence of IFN-γ for 48 hours, followed by GSIS experiments to see if GIP could diminish IFN-γ-induced functional impairment (Figure 2c). GIP neither affected baseline insulin secretion nor prevented the IFN-γ-mediated reduction in insulin secretion at low or high glucose (Figure 2e). We then tested if EndoC-βH5 cells exposed to IFN-γ for 48 hours retain the ability to respond to acute stimulation with GIP. In untreated control cells, 2.5 nM GIP, as expected, increased insulin secretion at both low and high glucose by 191% (*p*=0.06) and 164% (*p*=0.05), respectively (Figure 2f). In IFN-γ-exposed cells, acute GIP treatment appeared to increase insulin secretion at low and high glucose by 146% and 90%, respectively, although this was not statistically significant (both *p*=0.3).

**Figure 2.**
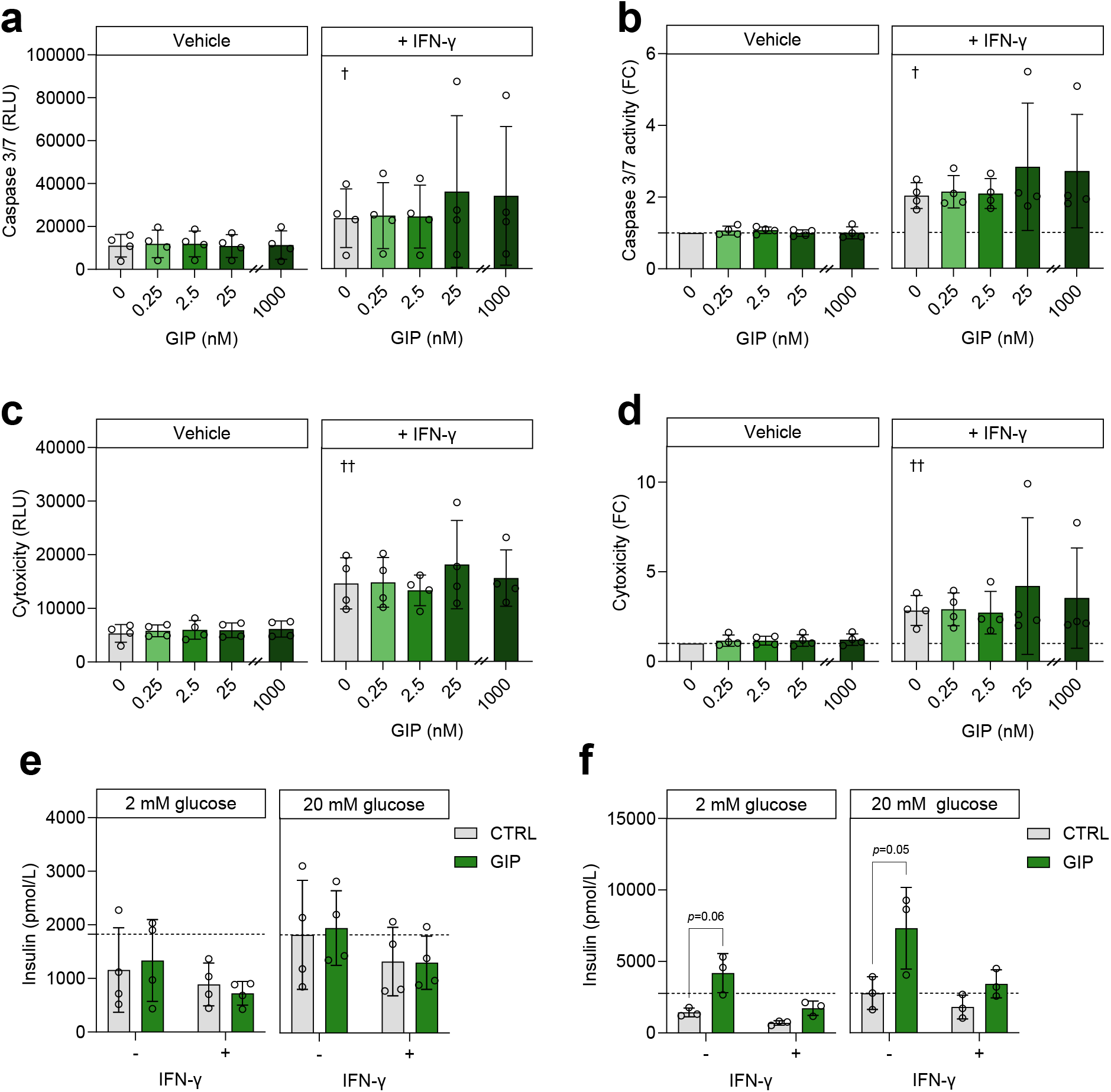
GIP does not prevent detrimental effects of IFN-γ exposure in EndoC-βH5 cells but cells tend to retain a partial response to acute GIP treatment after cytokine exposure. (a) Caspase 3/7 activity, (b) caspase 3/7 activity fold change, (c) cytotoxicity, and (d) cytotoxicity fold change in control (grey bars) or with increasing GIP (green bars, 0.25-1000 nM) after 96 h culture without (vehicle) or with IFN-γ (1000 U/mL). Angled lines on the x-axis denote the transition from x10 increments in GIP dose to 1000 nM GIP. Data presented as mean ± SD of 4 independent experiments. Non-significant using paired one-way ANOVA with Dunnett’s multiple comparisons test comparing each GIP dose to control (0 nM) at vehicle control or IFN-γ exposure, respectively. Daggers (†) indicate statistical significance using paired one-tailed Student’s t-tests comparing 0 nM GIP with or without IFN-γ. (e) Secreted insulin during low glucose (2 mM, left) and high glucose (20 mM, right) after 48 h culture without (grey bars) or with GIP (2.5 nM, green bars) in the absence (-) or presence (+) of IFN-γ (1000 U/mL). (f) After 48-hour culture in the absence (-) or presence (+) of IFN-γ (1000 U/mL) followed by acute GIP (2.5 nM, green bars) or not (grey bars). Data are represented as mean ± SD of 3-4 independent experiments. Non-significant using paired RM two-way ANOVA with Dunnett’s multiple comparisons test comparing GIP-treated to untreated controls in the absence of IFN-γ (-) or presence of IFN-γ (+) exposure, respectively. RLU, relative light units; FC, fold change.

### 3.4 GIP stimulates insulin and glucagon secretion from human islets but does not prevent cytokine-induced death of isolated human islets

To substantiate our findings on EndoC-βH5 cells, we evaluated the effects of GIP on human islets isolated from non-diabetic donors. At low glucose (2 mM, Figure 3a), there were no significant differences in the amount of secreted insulin upon treatment with 2.5 nM GIP compared to untreated control. At high glucose (20 mM), GIP-induced insulin secretion was 34.6% (*p*<0.001) higher than the untreated control, confirming an incretin effect. GIP significantly augmented glucagon secretion by 46.6% (*p*=0.01) at low glucose but not at high glucose (Figure 3b), confirming a glucose-dependent glucagonotropic effect of GIP. Consistent with our findings in EndoC-βH5 cells, GIP treatment alone did not affect baseline apoptosis, and co-culture with GIP failed to prevent apoptosis induced by IL-1β+IFN-γ exposure for 48 hours as evaluated by measurement of cytoplasmic nucleosomes using islets from 4 individual donors (Figure 3c-d).

**Figure 3.**
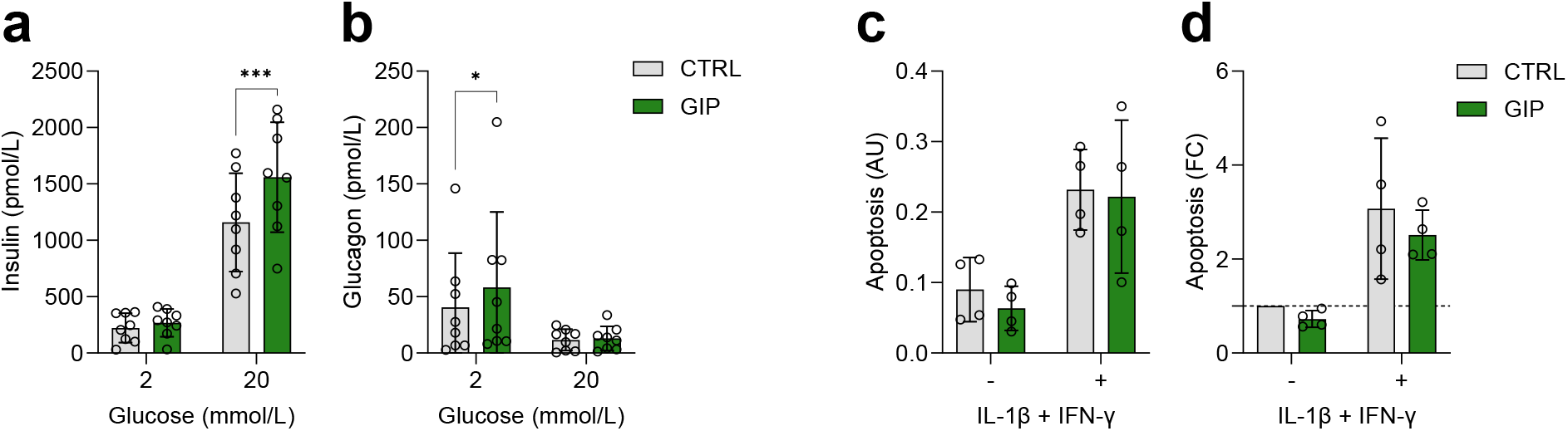
GIP is a dual-acting incretin hormone but does not prevent IL-1β+IFN-γ-induced apoptosis. (a) Insulin and (b) glucagon secretion at 2 and 20 mM glucose in control (grey bars) or with GIP (2.5 nM, green bars). Data presented as mean ± SD of 8 individual human islet donors. Asterisks (*) indicate statistical significance using paired RM two-way ANOVA with Šídák’s multiple comparisons test, comparing GIP-treated to untreated controls at 2 and 20 mM glucose, respectively, \**p* <0.05, \*\*\**p* <0.001. (c) Apoptosis (measured as cytoplasmic nucleosomes) and (d) apoptosis fold change in control (grey bars) or with GIP (2.5 nM, green bars) after 48 h culture with (+) or without (-) proinflammatory cytokine exposure (50 U/mL IL-1β and 1000 U/mL IFN-γ). Data presented as mean ± SD of 4 individual human islet donors. Non-significant using paired RM two-way ANOVA with Šídák’s multiple comparisons test, comparing GIP-treated to untreated controls without (-) or with (+) IL-1β and IFN-γ exposure, respectively. AU, arbitrary units; FC, fold change.

### 3.5 Cytokines, but not GIP, yield differential gene expression in human islets

Based on the lack of evidence for a cytoprotective effect of GIP in EndoC-βH5 cells and human islets, we sought to examine if GIP modulates the transcriptome of human islets, which could give insight into other potential functional effects, affected pathways and mechanisms. For this purpose, we performed RNA-sequencing (RNA-seq) of islets from 4 donors. Islets from each donor were cultured with or without 2.5 nM GIP in the presence or absence of IL-1β+IFN-γ for 24 hours. The RNA sequencing yielded an average of ∼46 million paired reads and ∼37 million aligned paired reads per sample (Figure 4a). We compared samples using Principal Component Analysis (PCA, Figure 4b) to visualise the treatment-induced transcriptomic changes. The PCA plot revealed that the greatest variance between samples (70% along PC1 on the x-axis) could be attributed to the proinflammatory cytokine exposure, as IL-1β+IFN-γ-treated samples clustered distinctively from non-L-1β+IFN-γ-treated samples. We also observed a 9% variance along PC2, where samples clustered according to the individual islet donors. Somewhat surprisingly, samples from GIP-treated islets appeared superimposed with their non-GIP-treated counterparts. Exploring additional PCs did not reveal GIP-directed clustering. Differential gene expression analysis of the 19,663 genes detected by RNA-seq revealed that the expression of 7,899 genes (∼40%) was significantly modified by the 24-hour exposure to cytokines with an equal distribution of up-and downregulated genes (Figure 4c and f). As eluted from the PCA plot, GIP treatment alone did not significantly alter gene expression (Figure 4d and f). Likewise, no genes were differentially expressed between the GIP+IL-1β+IFN-γ and IL-1β+IFN-γ groups (Figures 4e and f). IL-1β+IFN-γ treatment alone was robust and impacted both beta-cell and alpha-cell signature genes as defined previously [38] (Figure 4g and h) in line with the now-accepted view that also alpha cells are affected in type 1 diabetes [3]. Genes upregulated by IL-1β+IFN-γ treatment were enriched for processes related to immune and defence responses (Supplementary Figure 1). Interestingly, differentially expressed genes upon IL-1β+IFN-γ exposure were enriched at T1D risk loci; out of 152 pinpointed candidate risk genes from type 1 diabetes-associated loci [39], 96 were differentially expressed upon IL-1β+IFN-γ exposure (figure 4i), substantiating an interplay between islet inflammation and genetic risk. These findings indicate that while cytokines (as previously shown [40–42]) elicit profound changes to the transcriptome of human islets, GIP likely has minimal transcriptional impact. It is important to note that due to the limited sample size and the observed variability across donors, this data does not rule out a role of GIP in altering the transcriptome in human islets. Additionally, the response to GIP may exhibit more donor specificity, dosage dependence, and time-specific effects compared to cytokine treatment.

**Figure 4.**
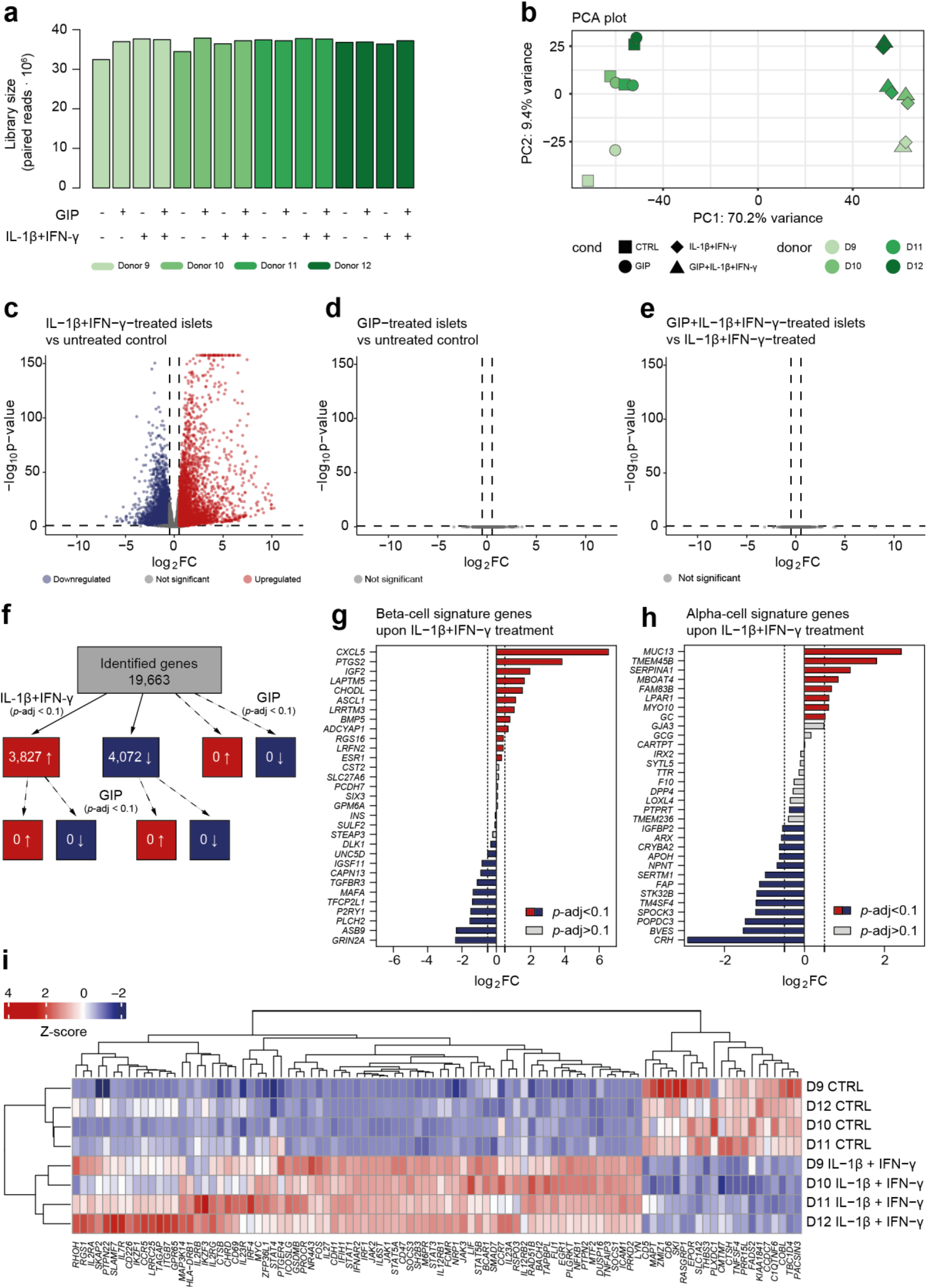
Cytokines, but not GIP, alter global gene transcription. (NEXT PAGE) (a) Library sizes for all 16 samples. (b) Principal component analysis of all 16 samples based on all expressed genes. (c) Volcano plot of differentially expressed genes in the IL-1β+IFN-γ-treated vs. untreated control contrast. Vertical lines denote log_2_ fold changes > |0.50|. Horizontal line denotes adjusted *p*-value < 0.1. Blue dots indicate significantly downregulated genes in the treated group, and red dots indicate significantly upregulated genes. (d) Same as in (c), but for the GIP-treated vs. untreated control contrast. (e) sample as (c-d) but for the GIP+IL-1β+IFN-γ-treated vs. IL-1β+IFN-γ-treated contrast. (f) Overview and number of differentially expressed genes in the three contrasts. Solid arrows denote IL-1β+IFN-γ treatment, while dashed arrows denote GIP treatment. (g) Impact of IL-1β+IFN-γ on beta-cell signature genes, and (h) alpha-cell signature genes. (i) Heatmap and hierarchical clustering of the 94 differentially expressed genes enriched at T1D risk loci.

### 3.6 qPCR validation of selected genes supports minimal transcriptional effects of GIP in human islets

To validate the RNA-seq findings, we performed qPCR of selected genes reported previously to be involved in GIP-mediated anti-apoptotic and survival effects in rodent beta-cell models); anti-apoptotic genes *BCL2* [15, 43] and *BCL2L1* (also known as *BCL-XL*) [17], pro-apoptotic genes *BCL2L11* (also known as *BIM/BIM-EL*) [17, 43] and *BAX* [17, 18, 43], and the endoplasmic reticulum stress marker *DDIT3* (also known as *CHOP*) [43, 44]. Consistent with the relative gene expression levels from RNA-seq, there were no differences in expression levels of the evaluated genes assessed by qPCR between GIP-treated and untreated control islets or between IL-1β+IFN-γ-treated and GIP+IL-1β+IFN-γ-exposed islets (Figure 5a-e). We furthermore performed qPCR of a few genes known to be induced by cytokines, i.e., chemokine-encoding genes *CXCL10* [6, 45, 46] and *CCL5* [47, 48], or *NOS2* [49]. Again, all three genes exhibited expression patterns comparable to the RNA-seq data (Figure 5f-h). Finally, we examined the impact of cytokines on *GIPR* expression by qPCR (Supplementary Figure 2). We observed a cytokine-induced ∼30% downregulation in the human islets, although the qPCR validation did not reach statistical significance (*p*=0.21).

**Figure 5.**
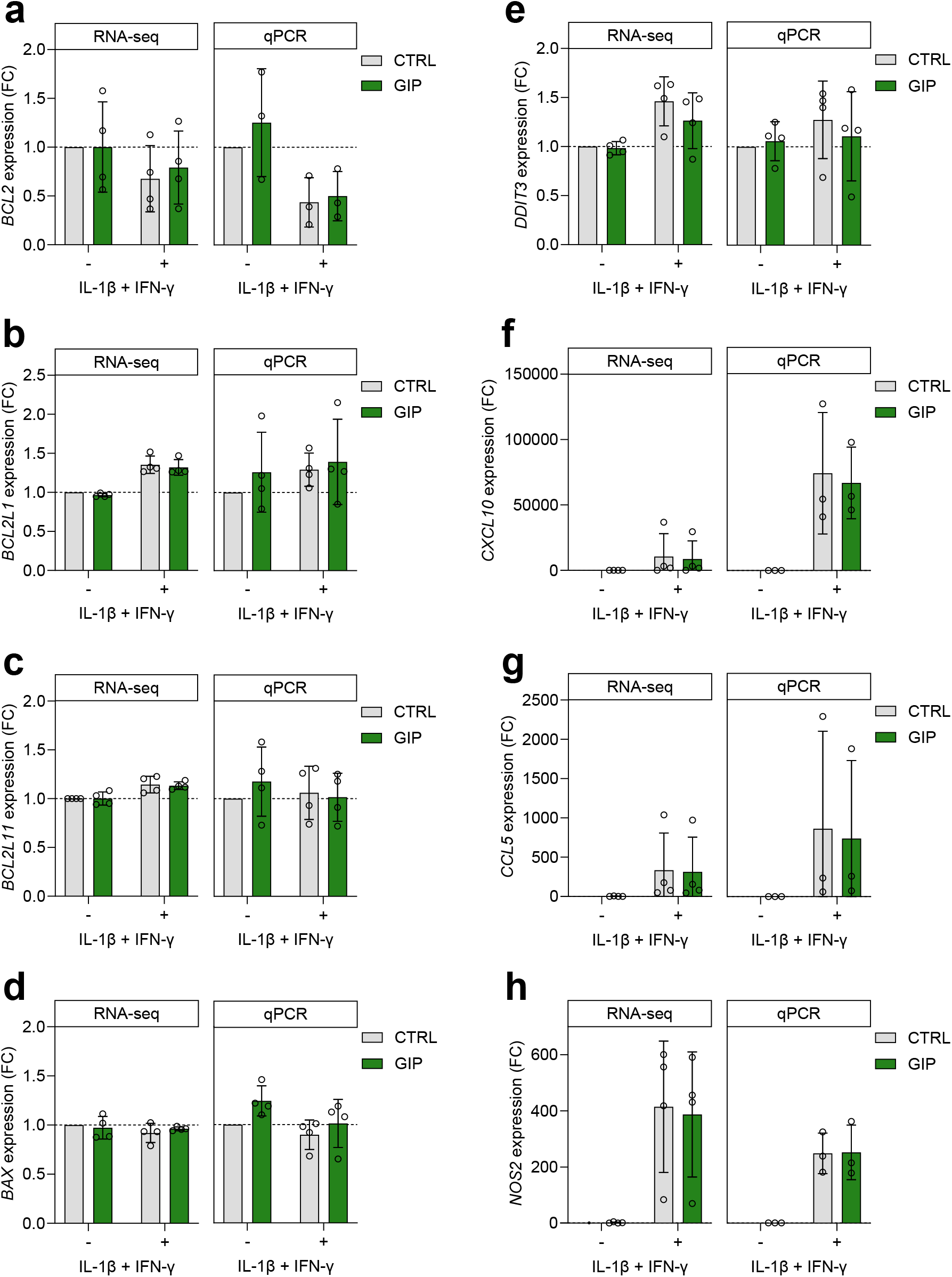
RT-qPCR validation supports RNA-seq gene expression. Gene expression of anti-apoptotic (a) *BCL2*, (b) *BCL2L1* (*BCL2-XL*), pro-apoptotic (c) *BCL2L11* (*BIM*), (d) *BAX*, (e) *DDIT3* (*CHOP*), and inflammatory markers (f) *CXCL10*, (g) *CCL5,* and (h) *NOS2* assessed by RNA-seq (right) or assessed by RT-qPCR (left). Data are represented as mean ± SD of 4 individual human islet donors (3-4 for qPCR). Non-significant determined by the DESeq2 Wald test for the RNA-seq data or paired RM two-way ANOVA with Šídák’s multiple comparisons test for the qPCR data, comparing GIP-treated to untreated controls without (-) or with (+) IL-1β and IFN-γ exposure, respectively.

### 3.7 GIP selectively modulates cytokine-induced secretion of inflammatory factors from human islets and EndoC-βH5 cells

Because islet cells secrete various chemokines and cytokines in response to pathophysiological stimuli [6–8], we measured the secretion of chemokines and cytokines from human islets exposed to IL-1β+IFN-γ ± GIP for 24 hours by using predesigned panels. Cytokine exposure significantly increased the secretion of all inflammatory factors measured, though not all reached statistical significance (Figure 6). GIP alone had no effects but significantly augmented cytokine-induced secretion of IL-10, CCL2, and CCL26 by 303% (*p*=0.03), 15% (*p*=0.006), and 45% (borderline significant, *p*=0.05), respectively (Figure 6e, -i, and -p). Further, GIP modestly but significantly decreased cytokine-induced secretion of CXCL8 by 11% (*p*<0.001, Figure 6g). We also measured the effects of GIP on the secretion of inflammatory factors from EndoC-βH5 cells (Figure 7) and found that GIP reduced IFN-γ-induced secretion of TNFα, IL-2, IL-6, and IL-10 by 29% (*p*=0.007), 17% (*p*=0.04), 17%, (*p*<0.001) and 31% (*p*=0.002), respectively (Figure 7a, -b, -d, and -e). In addition, GIP augmented IFN-γ-induced secretion of CXCL8, CCL2, CCL4, and CCL11 by 16% (*p*<0.001), 16% (*p*=0.005), 32% (*p*=0.04), and 36% (*p*=0.04), respectively (Figure 7g, -i, -k, and -l). These results suggest that GIP exerts selective inflammation-modulatory effects in human islets and EndoC-βH5 cells.

**Figure 6.**
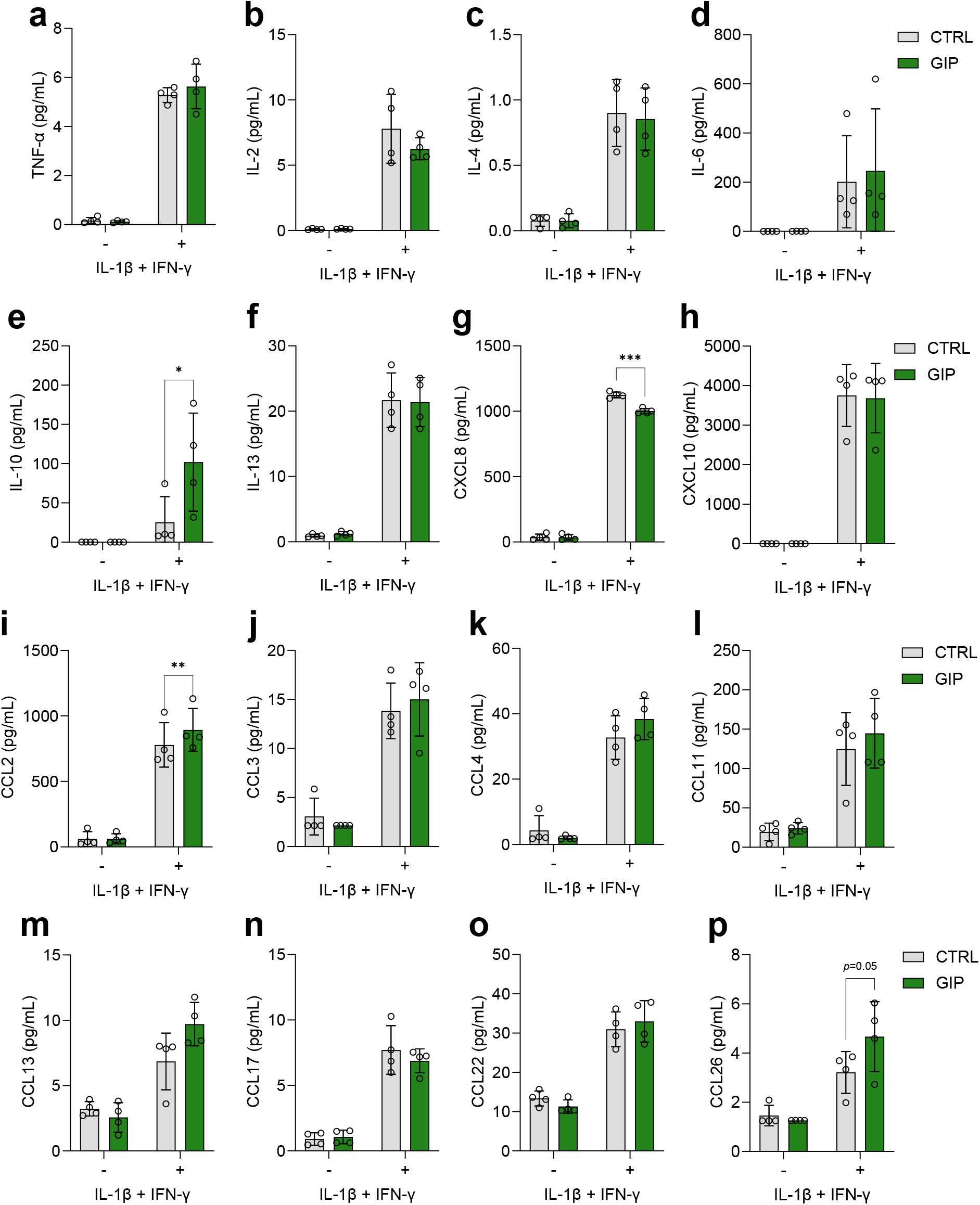
GIP treatment impacts cytokine-induced secretion of chemokines and cytokines in human islets. Accumulated secretion of (a) TNF-α, (b) IL-2, (c) IL-4, (d) IL-6, (e) IL-10, (f) IL-13, (g) CXCL8, (h) CXCL10, (i) CCL2, (j) CCL3, (k) CCL4, (l) CCL11, (m) CCL13, (n) CCL17, (o) CCL22, and (p) CCL26 during 24h of treatment without (grey bars) or with GIP (2.5 nM, green bars) in the presence (+) or absence (-) of proinflammatory cytokines (50 U/mL IL-1β and 1000 U/mL IFN-γ). Data presented as mean ± SD of 4 individual human islet donors. Asterisks (*) indicate statistical significance using paired RM two-way ANOVA with Šídák’s multiple comparisons test, comparing GIP-treated to untreated controls without (-) or with (+) IL-1β and IFN-γ exposure, respectively, \**p* <0.05, \*\**p* <0.01.

**Figure 7.**
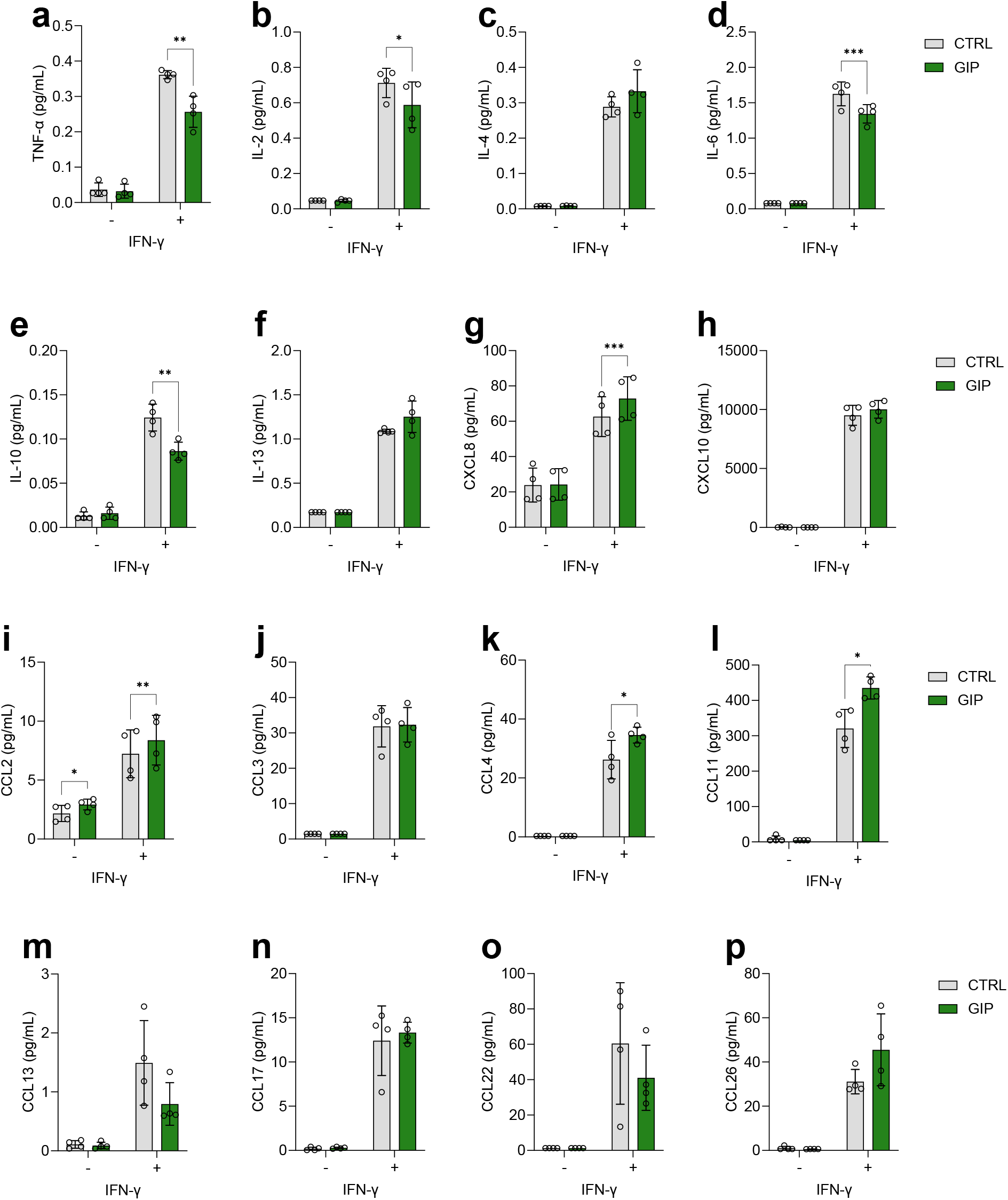
GIP treatment impacts cytokine-induced secretion of chemokines and cytokines in EndoC-βH5 cells. Accumulated secretion of (a) TNF-α, (b) IL-2, (c) IL-4, (d) IL-6, (e) IL-10, (f) IL-13, (g) CXCL8, (h) CXCL10, (i) CCL2, (j) CCL3, (k) CCL4, (l) CCL11, (m) CCL13, (n) CCL17, (o) CCL22, and (p) CCL26 during 24h of treatment without (grey bars) or with GIP (2.5 nM, green bars) in the presence (+) or absence (-) of 1000 U/mL IFN-γ. Data presented as mean ± SD of 4 independent experiments. Asterisks (*) indicate statistical significance using paired RM two-way ANOVA with Šídák’s multiple comparisons test, comparing GIP-treated to untreated controls without (-) or with (+) IL-1β and IFN-γ exposure, respectively, \**p* <0.05, \*\**p* <0.01, \*\*\**p* <0.001.

## 4. Discussion

To expand our insight into the effects of GIP on human islet endocrine cells, we explored the effects of GIP on EndoC-βH5 cells and human islets with a focus on insulin secretion, cell death, inflammation, and transcriptomic impact. Our findings suggest that (i) GIP-stimulated insulin secretion requires CaMK2, (ii) GIP likely has minimal protective effects against proinflammatory cytokine-induced beta-cell failure and death, (iii) GIP likely has minimal transcriptional impact in human islets, and (iv) GIP has selective modulatory effects on secreted inflammatory factors.

As recently reported for the newly established non-proliferative human beta-cell line EndoC-βH5 [26], we found that GIP acutely augmented insulin secretion at high glucose (20 mM) over a range of supraphysiological doses and modestly increased insulin secretion at low (2 mM) glucose. The increase in insulin secretion at low glucose may reflect that the tight regulation of beta-cell function is lost in 2D culture with a single islet cell type where paracrine signals from the delta and alpha cells are absent [50, 51]. This is further supported by the fact that we did not observe GIP-induced insulin secretion at low glucose in human islets in both this and a previous study [12].

CaMK2 has previously been shown to be involved in glucose-stimulated insulin secretion [52–54]. Consistent with this, we found that the CaMK2 inhibitor KN-93 abolished insulin secretion stimulated by high glucose alone. We furthermore found that KN-93 abrogated GIP-stimulated insulin secretion. Hence, both glucose-and GIP-induced insulin secretion from EndoC-βH5 cells appears to require CaMK2. In support, CaMK2 has been implicated in mitogenic and anti-apoptotic signalling by GIP in rat INS1-E cells [55], reinforcing the likelihood that CaMK2 could also be involved in GIP-mediated insulin release.

Ca^2+^ influx via LTCCs is a key event in the stimulus-secretion coupling process [56]. We found that the LTCC inhibitor nimodipine reduced insulin secretion, although this did not reach statistical significance (*p*=0.16) but did not affect GIP-induced augmentation of insulin secretion at high glucose, suggesting independence of Ca^2+^ influx via LTCC in EndoC-βH5 cells. As CaMK2 activity depends on Ca^2+^ binding to calmodulin (Ca^2+/^CaM), these findings indicate that GIP signalling may facilitate Ca^2+^ release from internal stores in human EndoC-βH5 cells, which should be further explored. The observation that PKA and PKC inhibitors failed to reduce GIP-stimulated insulin secretion at high glucose suggests a lack of dependency or redundancy for these components in GIP signalling in EndoC-βH5 cells. In support, there is evidence that the PKA inhibitor used, i.e., H-89, only partially diminishes GIP-induced insulin secretion in murine islets [57]. Redundancy in signalling might be achieved through other actors downstream of cAMP generation, such as EPAC2 [58]. EPAC2 may stimulate calcium release from internal stores via ryanodine and/or inositol 1,4,5-triphosphate receptors to induce insulin secretion [59]. To this end, it is unclear whether the discrepancies between previous studies and our study are due to technical and/or species differences. Therefore, additional experiments on human beta cells and islets should be performed to further decipher the signalling mechanisms of the hGIPR.

GIP failed to protect against cytokine-induced death of human EndoC-βH5 cells and human islets. Moreover, GIP did not prevent the IFN-γ-induced functional impairment, which precedes cell death of EndoC-βH5 cells [27]. These findings contrast earlier studies reporting cytoprotective effects of GIP in rodent islets and insulin-secreting cell lines [16, 18, 43, 60]. A single study used human islets and reported that 100 nM GIP partially counteracted cytokine-induced suppression of cell viability [61]. The study, however, evaluated viability using the MTS tetrazolium assay, which measures cellular metabolic activity as a representative measure of viability and thus may not provide sufficient information on the direct effect on cell death.

Recent evidence suggests that the human GIPR is more prone to internalisation and desensitisation in beta cells than the rodent GIPR [62, 63]. We, therefore, speculate if the lack of protective effects of GIP in our experiments with human cells could be due to reduced GIPR activity as a consequence of internalisation and desensitisation caused by the chronic GIP stimulation for 24-96 hours, resulting in insufficient GIPR signalling to exert cytoprotective effects. Alternatively, cytokines may reduce transcription of the *GIPR* gene. Assuming a congruence between mRNA and protein levels (yet to be confirmed due to the lack of specific antibodies for GIPR), cytokines may restrain GIPR protein expression and signalling in this way. Indeed, both in RNA-seq data from EndoC-βH5 cells (from [27]) and human islets, *GIPR* gene expression was downregulated by ∼30% in response to cytokines. Despite this, EndoC-βH5 cells partially retained the ability to respond to acute GIP stimulation following cytokine exposure, inferring that cytokine-induced reduced *GIPR* transcription is not the sole explanation for the lack of effect with GIP treatment alongside cytokine exposure. The lack of GIP protection against cytokine-induced functional impairment and cell death could, therefore, be due to a combination of *GIPR* downregulation and/or desensitisation and internalisation of GIPR. The latter is likely to have contributed more to the lack of GIP effect in the functional impairment experiments, as chronic GIP treatment did not increase insulin secretion in the absence of cytokine exposure.

Despite the inability of GIP to mitigate cytokine-induced cell death, it may still exert biologically relevant effects, potentially through gene expression modulation. However, global gene expression analysis of human islets revealed no significant alterations by GIP, either alone or upon cytokine exposure. This RNA-seq analysis was a 24-hour and single dose snapshot, and thus, potential transcriptional changes by GIP at earlier or later time points cannot be ruled out. Because GIP activates the transcription factor CREB in insulin-secreting rat INS-1 cells [15, 64] and other cell types [61], we expected CREB-dependent transcription, e.g., of proinsulin [65] or anti-apoptotic genes such as *BCL2* [15]. However, neither RNA-seq nor qPCR validation supported CREB-dependent transcription by GIP. To our knowledge, this is the first RNA-seq of human islets exposed to exogenous GIP. Few studies have investigated the impact of exogenous GIP in other tissues. One reported 911 differentially regulated genes in osteoblasts after a 4-hour treatment with GIP, but these were based on nominal p-values, not q-values or FDR [66]. In a clinical study with a six-day continuous subcutaneous infusion of GIP, no changes in global gene expression of white adipose tissue in type 1 diabetes individuals were found [67]. While we cannot definitively exclude that GIP impacts the human islet transcriptome, our findings suggest minimal or discrete transcriptional effects, possibly indicating a bias in GIPR signal transduction towards rapid and short-term posttranslational signalling mechanisms (such as CaMK2 activation) over slower transcriptional changes. Increased sample size may improve findings due to the substantial variance among deceased human islet donors. Since donors also likely show differences in cellular heterogeneity (alpha-beta-delta cell ratios), a single cell-level analysis would likely be very beneficial.

During type 1 diabetes development, the islets contribute to local inflammation by secreting various inflammatory factors that exacerbate immune cell infiltration and islet destruction [6–8]. GIP has been reported to have opposing impacts on chemokine and cytokine levels even within the same tissue or cell types [19]. In our data, GIP alone did not affect chemokine and cytokine secretion but strongly augmented IL-10 secretion from cytokine-exposed human islets. IL-10 is a cytokine with pleiotropic roles but was initially considered anti-inflammatory due to its actions as a negative feedback regulator [68, 69]. In human islet microtissues, co-treatment with IL-10 did not directly protect against cytokine-induced functional loss but reduced subsequent T-cell infiltration [70]. GIP-potentiation of IL-10 secretion from cytokine-stressed islets may dampen islet inflammation by reducing immune cell infiltration in situ. GIP also reduced cytokine-induced secretion of CXCL8 and increased secretion of chemokines CCL2 and CCL26, though to a much lesser extent than IL-While CCL2 promotes monocyte recruitment and islet inflammation in mice [71, 72] little is known of CCL26 in type 1 diabetes, but it may attract T-helper 2 (Th2) cells via the CCR3 receptor, which is also expressed by eosinophils and basophils [73]. Whether the increase in CCL2 and CCL26 outweighs the possible anti-inflammatory effects of augmented IL-10 secretion remains unclear at this point but deserves further investigation. Interestingly, GIP exerted a different modulatory effect on secreted inflammatory factors from EndoC-βH5 cells compared to human islets. Thus, GIP reduced IFN-γ-induced secretion of the cytokines TNFα, IL-2, IL-6, and IL-10, while increasing the secretion of the chemokines CXCL8, CCL2, CCL4, and CCL11. As for the human islets, whether the reductions in cytokines outweigh the increases in chemokines and how the overall inflammatory balance is affected warrant further research. The observed differences between human islets and EndoC-βH5 cells could potentially be due to cell type-specific responses to GIP, considering the presence of other endocrine cell types, primarily alpha and delta cells, in the islet. This possibility also merits further exploration.

In summary, our findings further our understanding of GIP biology at the human islet and beta-cell level and underscore the likely importance of investigating GIP effects in human cell models for valid translation to human physiology. In particular, our study warrants further investigation into the role of CaMK2 in GIP’s incretin effect and GIP’s inflammation-modulatory effects. Both will be important to further understand human GIP, its functional role, and the possible therapeutic potential in type 1 diabetes.

## 5. Additional information

## Acknowledgements

The authors gratefully acknowledge organ donors and/or their next of kin, without whom this research would not be possible.

## Data Availability

The RNA-seq data in this study are available in the Gene Expression Omnibus (GEO) repository under accession number GSE269993. The remaining raw functional data generated and analysed in this study are available from the corresponding author upon reasonable request.

## Contribution statement

KH and JS conceptualised the study and designed the experiments. AJ, RG, and CABS contributed to data acquisition. SK performed the initial prepossessing, mapping, and quantification of the RNA-sequencing data. KH analysed the data and drafted the manuscript. All authors critically revised the manuscript and approved the final version.

---

**Supplementary figure 1.**
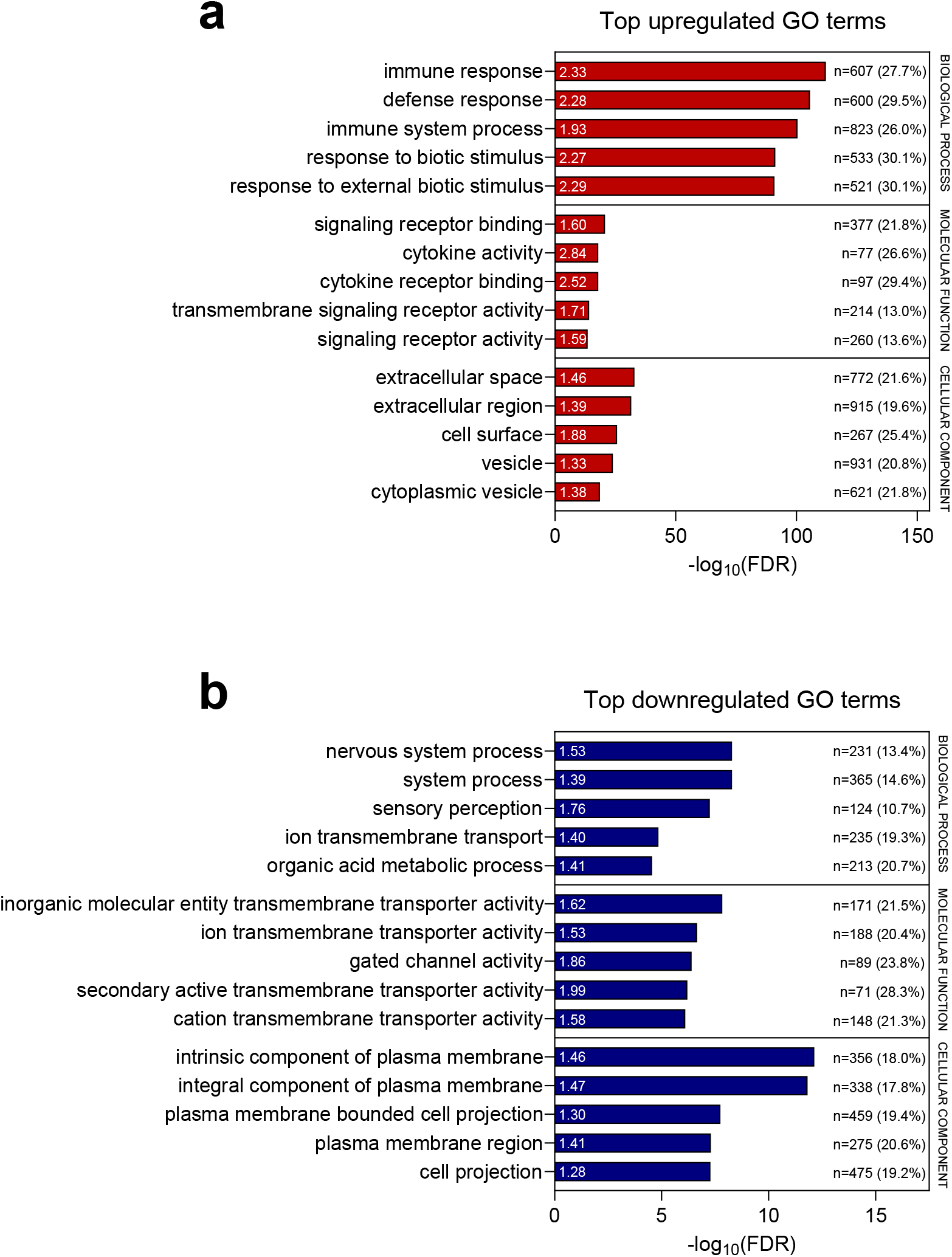
GO term enrichment upon IL-1β+IFN-γ. (a) Top 5 most significantly enriched GO terms (biological process, molecular function, and cellular component, respectively) of IL-1β+IFN-γ-upregulated genes or (b) downregulated genes. Fold enrichment of GO terms as indicated in white within each bar. The number (n) of regulated genes in the GO term and the percentage (%) of regulated genes out of the total genes in the GO term pathway as indicated in black adjacent to each bar. FDR, false discovery rate; GO, gene ontology.

**Supplementary figure 2.**
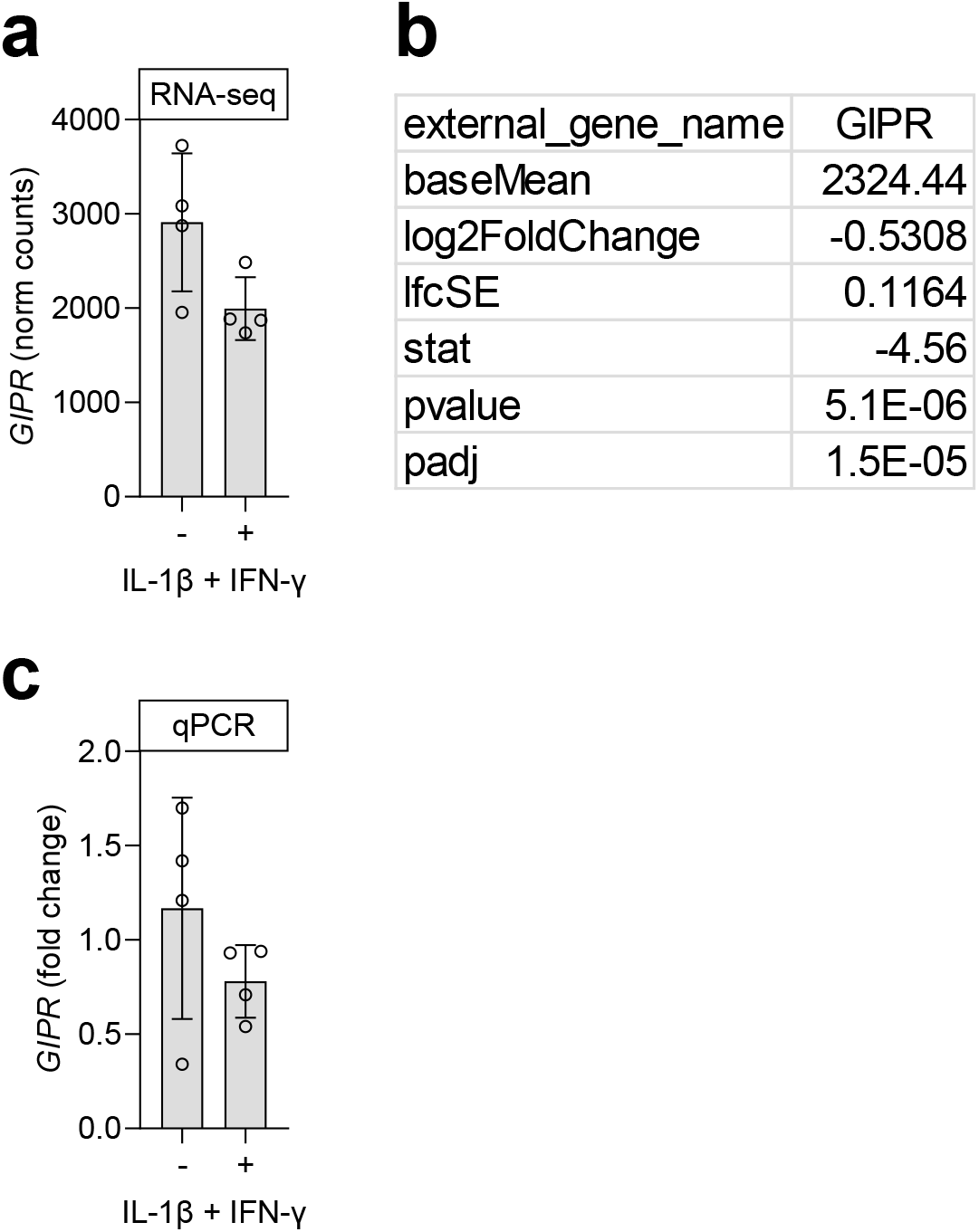
*GIPR* expression. (a) *GIPR* expression (normalised read counts) from RNA-seq and (b) the corresponding DESeq2 statistics. (c) *GIPR* expression evaluated by qPCR. Normalised to the average of controls to visualise spread of individual donors. Data are represented as mean ± SD of 4 individual human islet donors. Statistical significance determined by the DESeq2 Wald test for the RNA-seq (*p*<0.001) or two-tailed paired Student’s *t*-test for the qPCR data comparing untreated controls without (-) or with (+) IL-1β and IFN-γ exposure, respectively.

